# Multi Omics Driven Polypharmacology for Osteoarthritis Joint Regeneration

**DOI:** 10.1101/2025.08.13.670044

**Authors:** Amay Sanjay Redkar, Arun Surendran, Bishal Rajdev, Neethu Prasad, Arun N Prakash, Dikshita Hazarika, Ilackkeya Bhavananthi, Naveen Kumar, G Babu, YR Sanjayakumar, N Srikanth, Rabinarayan Acharya, Abdul Jaleel, VGM Naidu, CC Kartha, Sharad Pawar, Vibin Ramakrishnan

## Abstract

Osteoarthritis (OA) is a common degenerative condition of major joints with no effective cure. Yograj Guggulu (YG) is a well-established polyherbal formulation in Indian traditional medicine for the management of joint disorders. However, its mode of action remains unclear. We integrated multi omics network analysis with a rat model to investigate the effects of oral administration of YG on osteoarthritic joints. YG administration significantly alleviated pain and improved joint function. Imaging and histological analyses reveal that YG protects against structural degeneration in the joint with OA. Proteomic and metabolomic profiling, along with network analysis, uncovered that multiple components of YG alter protein expression in key pathways related to inflammation and cartilage degradation, thus validating the observed clinical effects. The efficacy of YG seems to be the result of the synergistic action of its diverse constituents. Our findings support the use of YG as a multitarget therapy for OA and encourage efforts for the potential repurposing of its active components.

## Introduction

Osteoarthritis (OA) is a degenerative disorder of major joints, eventually leading to disability, compromising the quality of life. Destruction of the articular cartilage, synovial membrane inflammation, and subchondral bone remodeling are the causes for the altered structure and function of joints in OA (Martel-Pelletier *et al*, 2016). The global prevalence of OA was 7.6 % of the population (595 million people) in 2020, a significant increase from 4.8 % (256 million) in 1990 (Steinmetz *et al*, 2023). Current pharmacological therapies are mainly palliative and do not cure or slow disease progression (Karsdal *et al*, 2016a). The most promising investigational drug is sprifermin, a recombinant form of fibroblast growth factor 18 (FGF-18) (Hochberg *et al*, 2019; Eckstein *et al*, 2021). This motivates the adoption of a mesoscopic paradigm, where disease modules arise from and are likely regulated by multi-level constraints (Bizzarri *et al*, 2011).

Yograj Guggulu (YG) is a polyherbal formulation used for centuries by practitioners of traditional Ayurveda to treat various degenerative conditions, especially joint disorders such as OA. It is prepared from a blend of 29 ingredients, with Guggulu (*Commiphora wightii* or *Commiphora mukul*) as the major component (Dutt *et al*, 2020). The different formulations and therapeutic uses of YG are extensively reviewed in *Bhaishajya Ratnavali*, an ancient Ayurvedic text. In the present study, we examined the possible mechanisms of action of YG on joints affected by OA.

Investigative approaches integrating proteomics, cheminformatics, and network medicine (Barabási *et al*, 2011) have provided a greater understanding of the pathogenesis and treatment of several complex diseases, such as cardiovascular disorders (Rai *et al*, 2021; Surendran *et al*, 2025; Lal *et al*, 2022), neurodegenerative diseases (Dou *et al*, 2025; Cummings *et al*, 2025; Varma *et al*, 2021; Seyfried *et al*, 2017), and cancer (Tan *et al*, 2019; Forbes *et al*, 2024; Yuan *et al*, 2021). We determined the efficacy and the molecular mechanisms of action of YG by integrating metabolomics, proteomics, and network analysis. We have further validated the results with imaging and histological studies of OA-affected joints in a Sprague Dawley rat OA model. Our results suggest that bioavailable metabolites from the polyherbal formulation YG, can impart disease-modifying effects in OA by modulating a set of topologically relevant proteins. This ayurvedic drug attenuates NF-κB signalling, extracellular-matrix degradation, and prostaglandin-driven inflammation that underpin joint degeneration.

## Results

### Disease Network Analysis

A network-based framework was adopted to identify the molecular mechanisms underlying OA, which involves the integration of genome-wide expression data (Figure 1, A). The topological analysis of this framework allowed us to identify key regulatory bottlenecks and control points. Further, integrative annotation and merging of data from previous studies were used to help position known OA markers. Ultimately, this framework has revealed the regulatory cascades in OA and the lead for recognizing the possible mechanisms of action of targeted therapeutic interventions.

**Figure 1:**
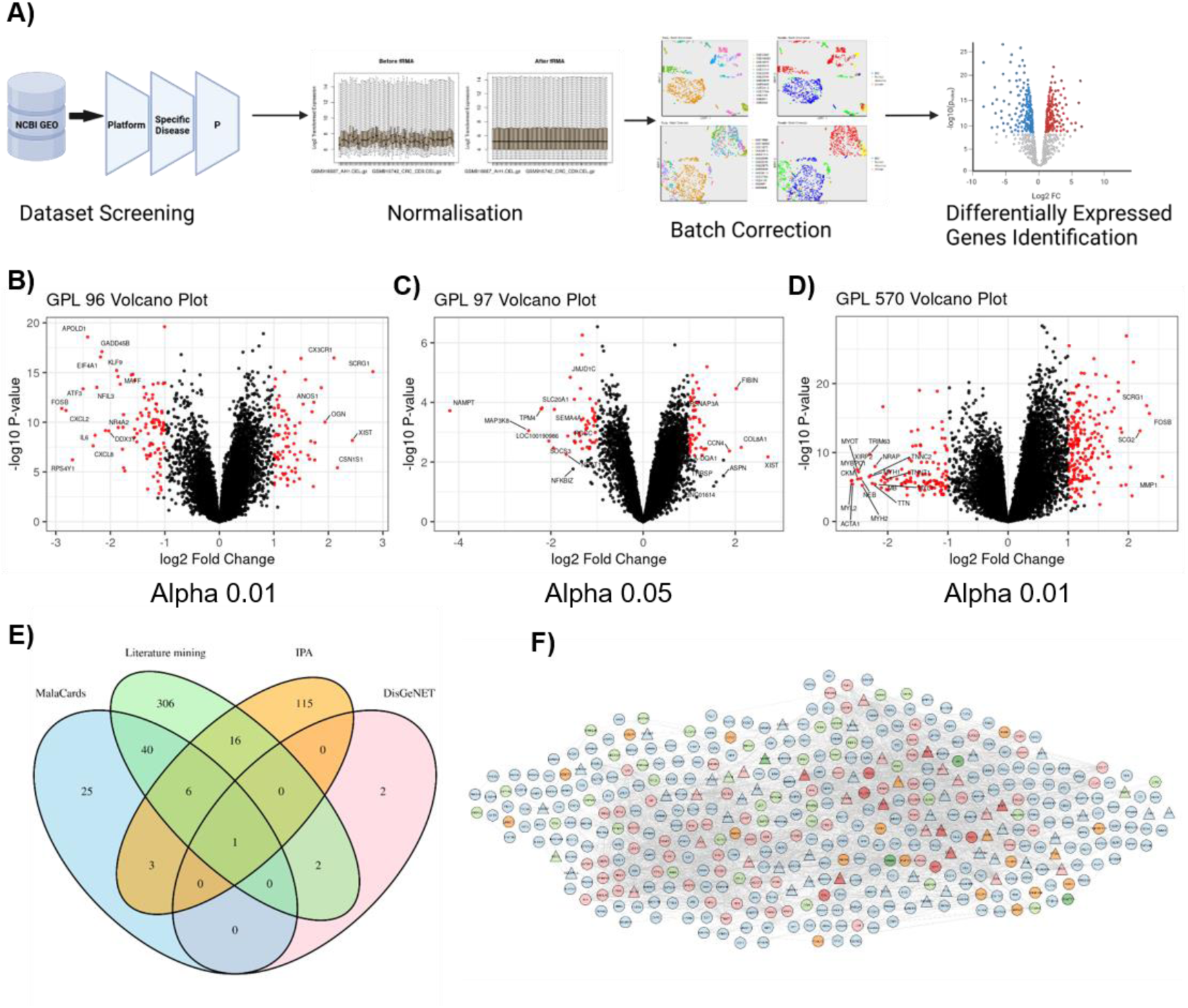
Overview of data processing, differential expression analysis, and OA disease network construction. (A) Schematic representation of the microarray data processing pipeline, from raw intensity files to normalized expression profiles. Created in BioRender. (B-D) Volcano plots of differentially expressed genes (DEGs) for the three platforms (GPL96, GPL97, and GPL570). Red points represent genes passing the chosen thresholds for statistical significance (y-axis) and fold change (x-axis). (E). Venn diagram showing the overlap of genes/proteins associated with OA from literature mining, DisGeNET, MalaCards, and IPA. (F) Protein-protein interaction network based on DEGs, with nodes representing individual genes and edges indicating inferred interactions. DEGs are colored light blue and round-shaped by default. Triangular nodes are genes that are drug targets. Hubs only, non-hub bottlenecks only, and OA associated only are colored light red, light green, and orange, respectively. OA-associated Hubs and OA-associated non-hub bottlenecks are colored dark red and dark green.

### Genome-wide Expression Studies

We have used a set of R (version 4.4.2) functions such as ReadAffy and fRMA. Each dataset was read, background corrected, frozen quantile normalized, and summarized using four R packages; affy (Gautier *et al*, 2004), fRMA (McCall *et al*, 2010, 2011b, 2011a), hgu133afrmavecs, and hgu133plusfrmavecs. The specific usage of these tools depended on the platform and series multiplicity. Following pre-processing and normalization, ten Gene Expression Omnibus (GEO) datasets were merged by the platform. They were collectively assessed as three datasets with 295 OA and 57 normal synovium samples. The GPL96 platforms had four series with 65 samples (29 normal, and 36 OA). The GPL97 platform included one series of 14 samples (4 normal and 10 OA). The GPL570 platform had five series of 273 samples (24 normal and 249 OA). Dimensionality reduction was performed using Uniform Manifold Approximation and Projection (UMAP) (McInnes *et al*, 2018). The batch effects were identified and subsequently eliminated by empirical Bayes estimation, or ComBat from sva (Leek & Storey, 2007; Leek *et al*, 2019). This correction effectively removed the non-biological clustering, thus facilitating the observation of only the biologically relevant differences (Figure S1).

Differential expression analysis identified 544 genes as differentially expressed (DEGs) (Data S1 to S5) in the OA compared to normal samples, based on a threshold of |LogFC| ≥ 1.0 and FDR q-value < 0.01 for integrated datasets. For the analysis of a single dataset, while keeping a threshold of |LogFC| ≥ 1.0, a less stringent FDR q-value < 0.05 was used (Figure 1, B to D, table S1). LogFC and FDR refer to Log [Fold Change] and False Discovery Rate, respectively.

### Network Construction

The DEGs identified in OA represent a promising set of potential seed nodes that can be involved in the development of OA or at least in the presentation of its symptoms in humans. These genes can be involved in the disease phenotypes, potential therapeutic targets, or both. These targets could control the network’s topological structure and the phenotypic state of OA. Integrating these genes into the network might enable us to discover key regulatory elements and pathways involved in the progression of OA.

We constructed undirected and directed networks using nodes derived from integrated DEGs as seeds to elucidate the system-level interactions in OA. The undirected networks were constructed using an adjacency matrix based on protein correlation and co-occurrence data from the STRING database (Szklarczyk *et al*, 2023). Directed networks were assembled using interaction data from NetControl4BioMed (Popescu *et al*, 2021), which complies with experimentally validated information from Omnipath (Türei *et al*, 2016), InnateDB (Breuer *et al*, 2013), and SIGNOR (Licata *et al*, 2020) databases to pinpoint nodes that modulate disease control. Additionally, these networks were expanded by incorporating the neighboring nodes of the DEGs to gain deeper insight into the mechanism of the activated pathway (Data S6). This extension is based on the premise that the DEGs are mainly indicators of disease-altered processes, rather than the “causes” of the disease at the transcriptome level (Varma *et al*, 2021; Porcu *et al*, 2021). Hence, the control nodes (disease-associated and druggable) identified within the expanded network may hold key regulators that are not differentially expressed, but could be crucial in orchestrating the phenotypic changes that characterize the disease, thus potentially revealing novel potential therapeutic targets (Vinayagam *et al*, 2016; Nogales *et al*, 2022; Rheinbay *et al*, 2020).

Following the network construction, topological analysis was employed to explore the interaction between nodes. This involved mapping of protein-protein interactions (PPIs) as well as examining the properties of the networks as a whole (Table S2). We have identified the key hubs (high-degree nodes) and the non-hub bottlenecks (nodes with high betweenness but low degrees) in the network’s topological properties, collectively called Topologically Relevant Genes or TRGs (Figure S2, table S3). This model offers a conceptual framework to understand how critical interactions might contribute to pathological changes in OA, as well as identify specific nodes that can alter the course of OA when manipulated.

### Network annotation

Initially, we identified a set of 516 genes and proteins associated with OA. From this set, 371 genes/proteins with a p-value of ≤ 0.05 were identified through literature mining. Specifically, five genes/proteins were sourced from DisGeNET (Piñero *et al*, 2020), 75 from MalaCards (Rappaport *et al*, 2017), and 141 were from IPA (Data S7). To ensure complete coverage of OA-associated biomarkers, all the identified genes/proteins were included for further investigation, regardless of their frequency in the databases (Figure 1, E). Additionally, known drug targets were extracted from the DrugBank database (Knox *et al*, 2024) to integrate therapeutically relevant information into the network (Data S8).

### In vivo validation of the effect of treatments against OA

We used the OA disease network, which is annotated with known disease-associated genes, drug targets, and network-controlling nodes, to test the therapeutic efficacy of YG. Our predictions were validated in a monoiodoacetate (MIA)-induced OA model in Sprague Dawley rats. MIA, a metabolic poison, blocks glyceraldehyde-3-phosphate dehydrogenase (GAPDH), inhibiting the glycolytic pathway and induces chondrocyte apoptosis. Chondrocyte death leads to extensive neovascularization, subchondral bone necrosis and collapse, prolonged inflammation, osteophyte formation, and degradation of articular cartilage (Aüllo-Rasser *et al*, 2020).

The disease control (DC) group and the treatment groups, low (YL), medium (YM), and high (YH) doses of YG and diclofenac sodium (STD), received an injection of MIA into the knee joint of the left hind leg. To account for potential confounding effects from needle-induced damage to small structures of the rat knee joint, a vehicle control (VC) group was also included. Given that there are no approved disease-modifying treatments for OA (Karsdal *et al*, 2016b)Diclofenac sodium, an anti-inflammatory analgesic, was used as the standard control.

An injection of MIA into the knee joint induced OA on day one. After seven days, oral administration of YG or diclofenac sodium was continued up to 21 days (Figure 2, A and B).

**Figure 2:**
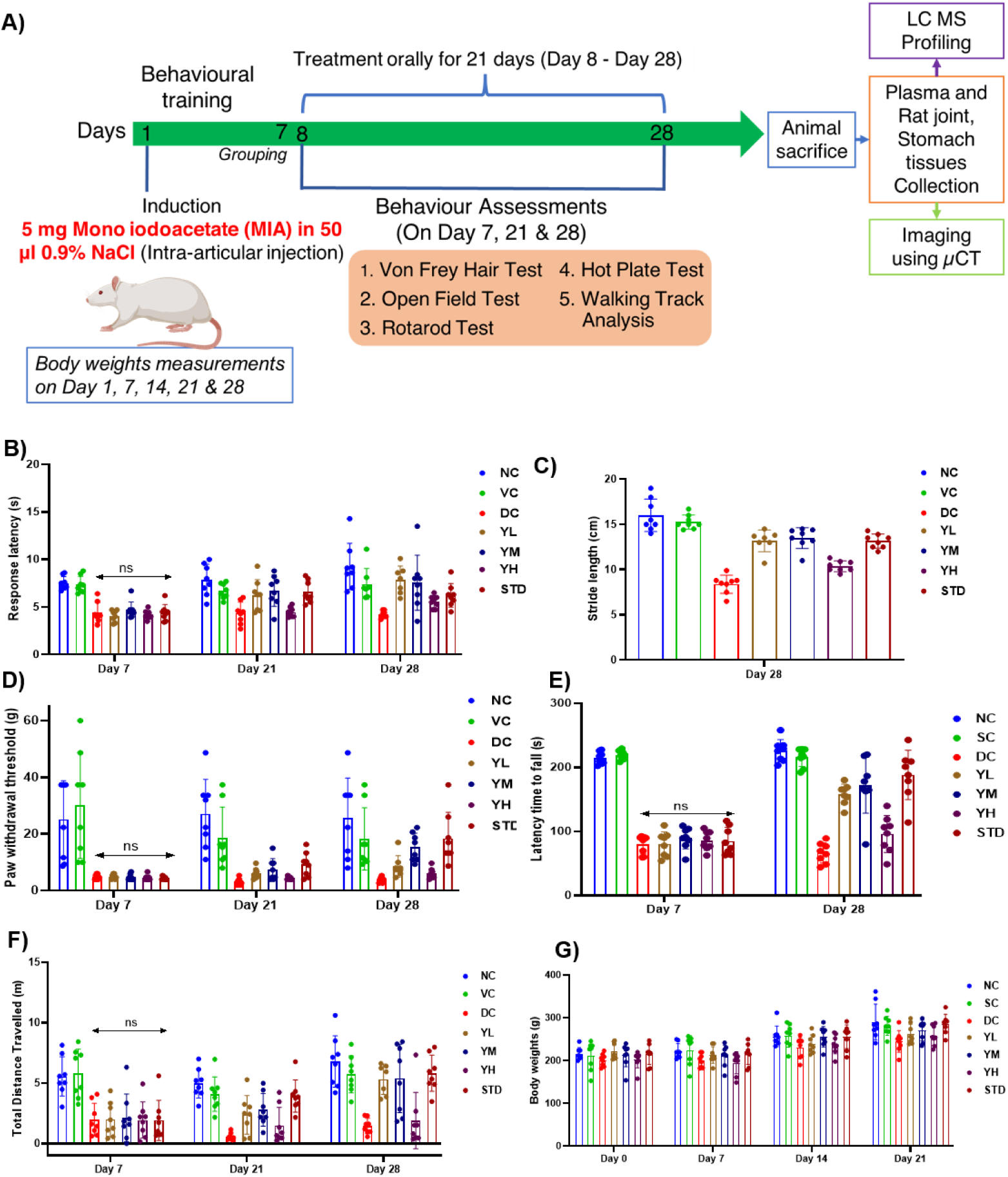
Experimental design, treatment regimen, and behavioral outcomes in an MIA-induced rat model of Osteoarthritis. (A) Schematic representation of the study timeline. Rats were injected with monosodium iodoacetate (MIA; 5 mg) on Day 0 to induce osteoarthritis-like changes. Following 7 days of behavioral training (Days −7 to 0), oral treatments were administered daily for 21 days (Days 8–28). During this period, various behavioral tests (Von Frey, Open Field, Rotarod, Hot Plate, and Walking Track) were conducted to assess pain sensitivity and locomotor activity. At the end of the treatment, animals were sacrificed for tissue collection and further analysis (stomach and knee joint histopathology, micro CT of the knee joint, plasma metabolomics, and knee joint proteomics). Created in BioRender. (B–G) Bar plots show the effects of different treatment groups on the behavioral outcomes at multiple time points. Error bars represent mean ± SD (n = number of rats per group). Significant differences, determined by two-way ANOVA followed by Tukey’s post hoc analysis, between groups and time points highlight the efficacy of each treatment in mitigating osteoarthritis-associated pain and mobility deficits.

### Efficacy of YG based on behavioral studies

Seven days after the induction of OA using MIA, disease induction was verified using behavioral tests, including the hot plate, rotarod, open field, walking track, and Von Frey hair tests (Data S9 to S13). These tests were used to measure pain response to thermal stimuli, motor coordination and balance, general locomotor activity, and tactile allodynia. All tests confirmed significant impairment of knee joint function in the affected groups, indicating successful induction of OA-like conditions in the rats.

After 14 days of treatment with YG and diclofenac sodium (21 days post-MIA injection), joint functions were reassessed using the hot plate, open field, and Von Frey hair tests. We observed locomotor activity was enhanced in the treated groups YL, YM, and the Standard Control. The YH group also showed improvements, but not to the same extent as in the other groups. There was an enhancement in thermal nociceptive response, measured by the increased hot plate response latency in the YL, YM, and standard control groups (Figure 2, B to G). Pain sensitivity assessed by measuring the paw withdrawal threshold was unchanged in the treatment groups compared to the DC group.

Reduced paw print distances because of OA-related changes in gait patterns were observed in the DC group after 21 days. Conversely, the gait patterns revealed signs of recovery in the YL and YM groups. In the hot-plate test, response latency improved on the 28^th^ day of treatment. The YL group demonstrated the most significant reduction in nociception, followed by the YM group. Compared to the DC group, the standard group showed the most substantial recovery in locomotor activity, followed closely by YM and YL. These findings collectively highlight the efficacy of our treatment regimen in restoring joint movements.

The improvement in the paw withdrawal threshold in the YM group, compared to the DC group, suggests alleviation of tactile allodynia on treatment. The recovery observed in the treatment groups was similar to that observed in the NC group. There was no change in body weights of the animals during the study, indicating that fluctuations in body weight did not influence the changes in the mobility and pain perception (Figure 2, H, Data S14).

Our results indicate that oral administration of YG or diclofenac sodium effectively ameliorated the OA-induced dysfunction with notable improvements in pain sensitivity, locomotor activity, and gait patterns.

### Efficacy of YG assessed by micro-CT imaging of the knee joint

Three weeks after YG administration, the knee joints of the rats were evaluated using micro-CT imaging. Our findings indicate that both YG and diclofenac sodium caused improvement in multiple parameters (Figure 3, A, Data S15). Specifically, the tibial bone mineral density (BMD) was lower in the DC group compared to the NC group. Tibia’s BMD was moderately higher in the SC, YM, and YL groups than in the DC group (Figure S3, A). Fibula’s BMD was very low in the DC group and moderately high in the SC, YM, and YL groups (Figure S3, B). Both the diclofenac sodium and YG-treated groups showed increased bone volume compared to the disease control group, suggesting an improvement in bone mass (Figure S3, C) and mitigation of bone degradation, typically observed in OA. There was also restoration of cortex volume to different degrees in the treatment groups (Figure S3, D and E). Improved cortical volume correlates with fewer bone-marrow lesions (a strong pain driver) and slower cartilage loss. The treatment groups showed an improvement in the trabecular tissue volume (Figure S3, F to H).

**Figure 3:**
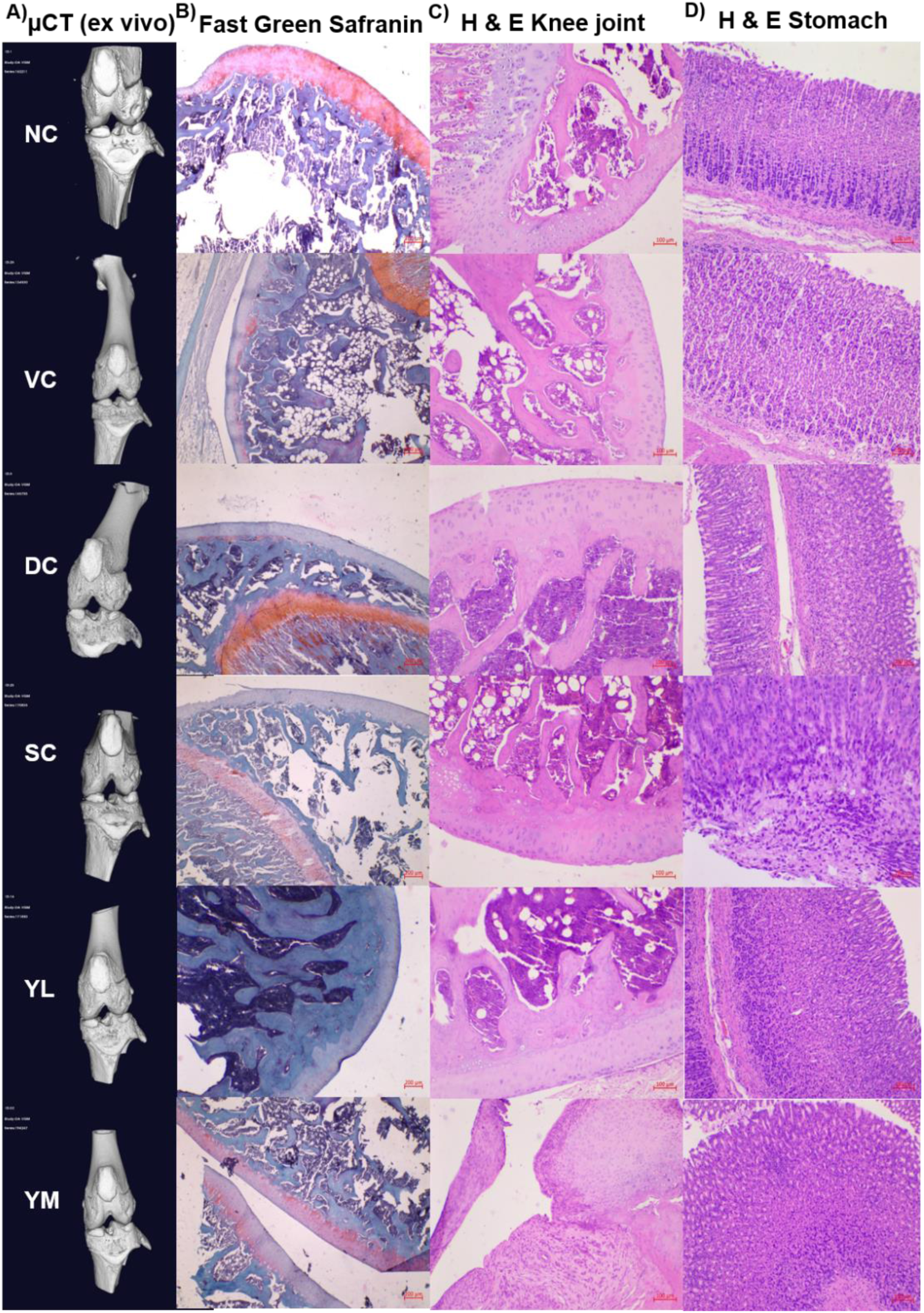
Micro-CT and histological assessments of the effects of treatment MIA-induced OA rat knee joint and stomach. (A) Representative micro-CT reconstructions of the knee joint, illustrating structural changes in the femoral condyle and tibial plateau across different treatment groups. (B) Representative Safranin O/fast green stained sections highlighting proteoglycan loss (reduced red staining) in OA-affected joints compared to normal controls. (C) Representative Hematoxylin & eosin (H&E) stained sections showing cartilage erosion, chondrocyte organization, and synovial changes. (D) Representative Hematoxylin & eosin (H&E) stained sections showing changes in the stomach’s morphological structure and mucosal lining.

The betterment observed in imaging parameters indicated the efficacy of YG in maintaining trabecular architecture, suggesting a restorative effect against trabecular and bone loss, which are often associated with OA.

### Efficacy of YG was assessed based on the histopathology of the knee joint

Three weeks after YG administration, knee joint tissues were harvested from rats and processed for histology. Cartilage degradation was evaluated using a grading system of Osteoarthritis Research Society International (OARSI), which is based on Fast Green Safranin staining of the tissues (Figure 3, B, Figure S3, I). This method enables a detailed assessment of the proteoglycan content, chondrocyte viability, and matrix organization within the articular cartilage. In this grading system, the intensity of the red stain reflects the proteoglycan levels, while changes in the cellular arrangement and the presence of fissures, erosion, and fibrocartilaginous remodeling indicate progressive cartilage degeneration.

The NC group displayed intact morphological features of the cartilage, including uniformly arranged chondrocytes, robust proteoglycan content, and structurally preserved trabecular bone (Grade 0). The VC group also presented relatively normal morphological features, with mild degenerative changes observed in two samples. In contrast, the MIA-induced OA DC group showed severe cartilage erosion, matrix depletion, complete loss of chondrocytes, as well as proteoglycan depletion, and marked inflammation were consistently observed (Grade 4–6). Significant protective effects were seen with the YL and YM doses. Significant focal degeneration, cartilage erosion, and inflammation were seen in rats that received high doses of YG, highlighting potential risks of high-dose intake. Animals that received the standard drug, diclofenac sodium (STD), maintained normal cartilage structure. Although mild focal degeneration and inflammatory reactions were occasionally noted.

Histological features of the knee joints are shown in Fig. 3C. The NC and most VC animals showed normal articular cartilage with normally aligned chondrocytes, intact extracellular matrix, normal trabecular bone, and a typically normal synovial membrane. On the other hand, the DC group displayed typical osteoarthritic changes, such as thinning of the matrix, loss of chondrocytes, and multiple focal areas of bone resorption, accompanied by inflammatory pannus formation. Animals in the YL and YM groups had near-normal cartilage architecture, while the YH group presented severe erosive and inflammatory damage. The standard treatment group (diclofenac sodium, STD) demonstrated only partial recovery, with cartilage loss and inflammatory changes similar to those observed in the DC group.

Taken together, these observations suggest that YG has protective effects on cartilage with varying efficacy depending on the dose.

### Toxicity evaluation of YG based on histopathology of the stomach

Histology of the gastric wall revealed distinct features in different animal groups (Figure 3, D). The NC and VC groups had no abnormalities of the stomach’s mucosal, submucosal, and muscular layers. The DC group had multifocal mild inflammation and lymphocytic infiltration, predominantly in the mucosa and muscularis mucosa. In the YL, YM, and YH groups treated with YG, the gastric wall was mostly normal. A few specimens had mucosal erosion, focal inflammatory infiltrates, and fibrous tissue proliferation. Pathological changes such as extensive ulceration, granulation tissue formation, mixed inflammatory cells, and focal necrosis, indicative of severe tissue damage, were seen in the STD group.

Histopathology thus reveals that the diclofenac sodium intake over a long term can lead to severe gastric mucosal damage. YG seems a safer option for long-term intake.

### Chemical Constituents, Drug-like Compounds in YG and their targets

After 21 days of treatment with YG, blood was drawn from MIA-induced OA rats by cardiac puncture. The full range of YG-induced metabolic changes is unknown. We used untargeted metabolomics to get an unbiased snapshot of all plasma metabolites altered by the treatment. This allows us to detect all bioactive bioavailable compounds. Untargeted metabolomics of the blood plasma was conducted to identify metabolites, annotated using Compound Discoverer, that are differentially abundant in YL-treated MIA-induced OA rats compared to the untreated MIA-induced OA rats and to identify structurally similar entities from YG (Data S16 to S21). Differential abundance analysis of Compound Discoverer identified 98 metabolites as differentially abundant metabolites (DAMs) in the YL versus DC comparison (Figure S4). Among these, 47 unique, similar drugs with more than 0.6 Tanimoto coefficient similarity were identified. The Tanimoto coefficient compares two molecular fingerprints, and a score ≥ 0.6 is widely accepted as evidence of meaningful structural similarity that can hint at shared bioactivity. These drugs are associated with 34 known unique targets (Figure 4).

**Figure 4:**
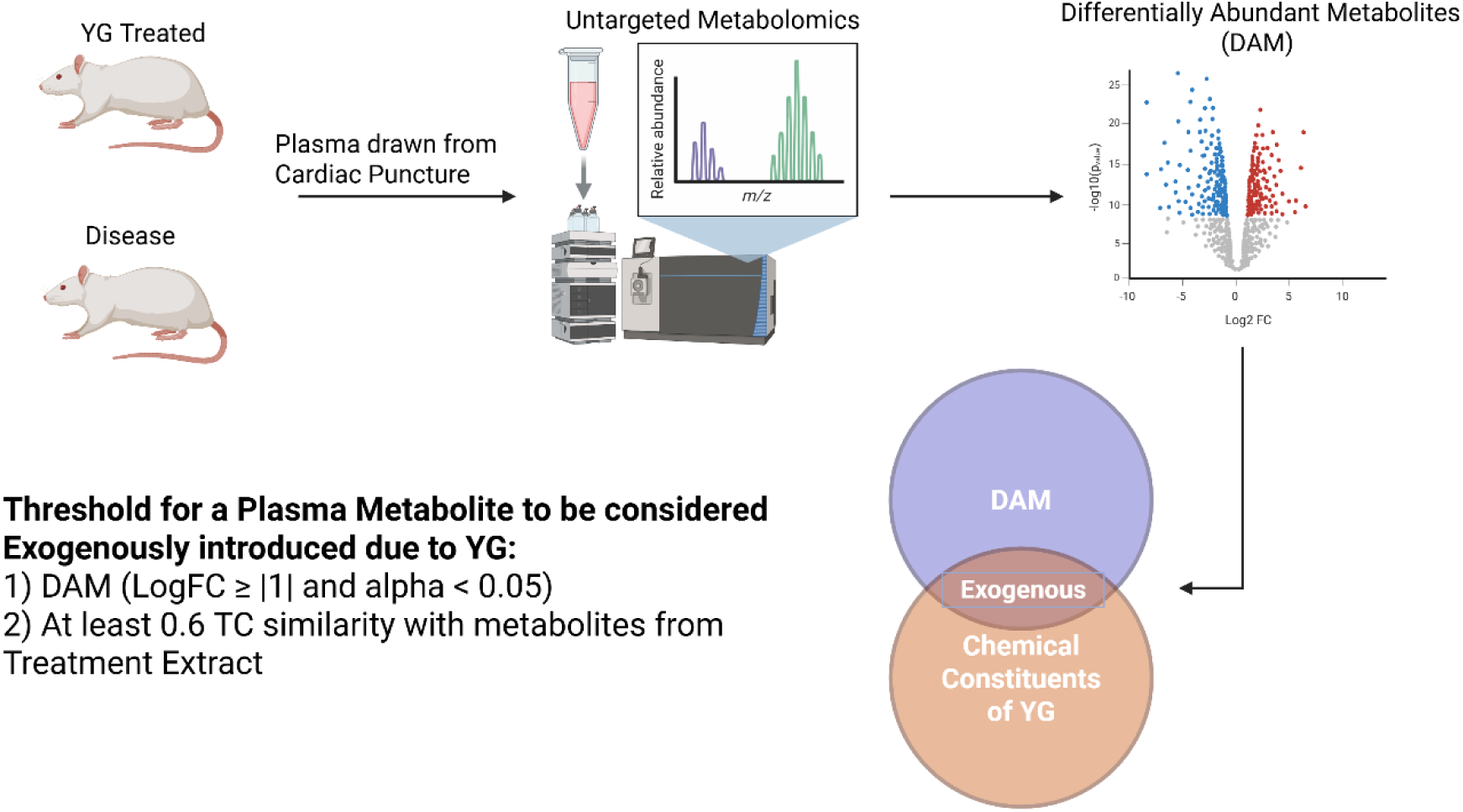
Untargeted metabolomics workflow, identification of DAMs, and multivariate analysis of plasma samples. Schematic illustration of the overall metabolomics approach, from sample collection in control and treated rats to LC-MS-based analysis and data processing. Created in BioRender. The summary table S5 lists the treatment groups, representative plasma metabolites, and exogenous compounds with similar drug structures and their corresponding target coverage.

To identify metabolites that YG exogenously introduced, we profiled the methanol extract (20 mg of YG in 1 ml 80 % chilled methanol with 0.1 % formic acid) using untargeted metabolomics (Data S22 to S30). The identified compounds were then compared for their structural similarity to the DAMs in the YL versus DC comparison to determine exogenously introduced metabolites based on the structure-activity relationship. DAMs with at least a Tanimoto coefficient similarity value of 0.6, to those from the extract, were considered exogenously introduced. Of these, 23 YG metabolites showed similar bioavailable metabolites and were found to have 289 unique drug targets, which were identified based on the structure-function similarity principles (Table S5).

### Proteomics Analysis

To examine the therapeutic effect of YG at the molecular level and extrapolate its functional changes, we conducted untargeted proteome profiling of the knee joint tissue of rats with MIA-induced OA (Figure 5A). For this profiling, we selected tissue samples from the YL group (one of the three YG doses used), as well as from the normal (NC), disease (DC), vehicle (VC), and diclofenac sodium treatment (STD) groups. Four samples from each group were analyzed with each sample profiled in triplicate (Figure 5B).

**Figure 5:**
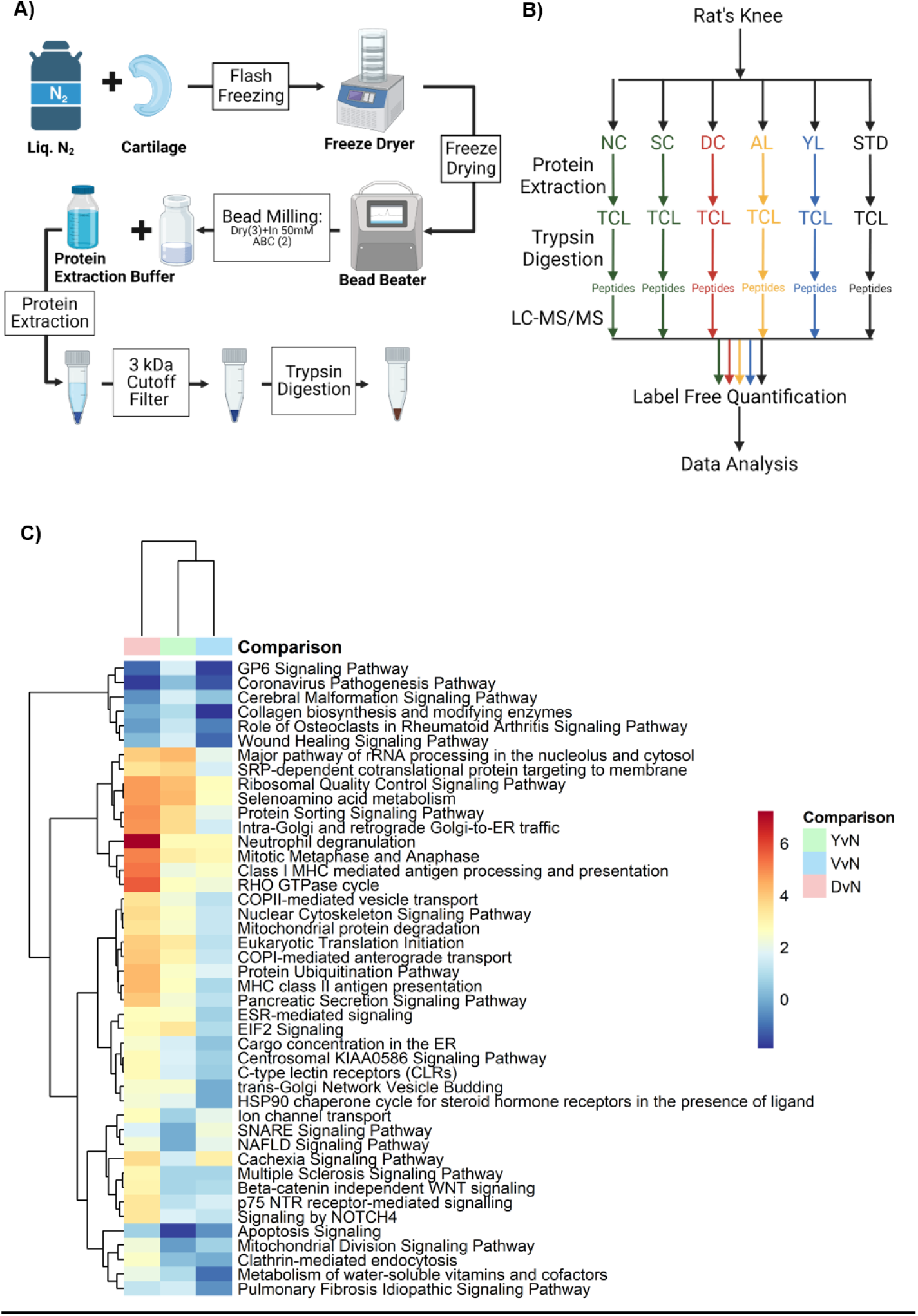
Proteomic workflow, PCA-based classification of knee joint samples, and functional enrichment analysis of differentially expressed proteins. (A) Schematic representation of the proteomic pipeline, from flash-freezing and bead milling of knee joint tissue through protein extraction, size exclusion (3 kDa cutoff), and trypsin digestion. Created in BioRender. (B) Experimental grouping and treatment scheme, illustrating how the treatment and controls were processed for downstream analyses. Table S6 summarizes the DAPs in treatment groups compared to normal control. (C) Comparison canonical pathway analysis using IPA of the DAPs from DvN, YvN and VvN groups as heatmap. The pathways related to OA are activated in DvN but not in YvN and VvN highlighting YG’s ability to dampen OA-associated inflammatory signalling and partially restore tissue-repair pathways.

Total Peptide Amount (TPA) normalization was used to normalize the samples. The TPA normalization is a method that groups all the peptides in each sample, calculates the total signal for all the samples, and then selects the maximum signal. The normalization factor was calculated for each sample by dividing the peptide sum of that sample by the maximum peptide sum across all the samples. This approach effectively corrected for most variations in total peptide yield, allowing for a comparative analysis of different conditions (Figure S5).

The variance of the dataset was evaluated using Principal Component Analysis (PCA). PC1 and PC2 account for 25.4 % and 12.4 % of the rat knee joint proteomic variance respectively (Figure S6). The samples were separated into two main clusters along PC1, distinguishing healthy (normal control) from MIA-induced OA samples. This separation highlights significant differences in protein expression profiles that exist between the groups, which align with our findings from behavioral, micro-CT, and histopathological experiments. Moreover, within the MIA-induced OA group, samples were further dispersed along PC2, suggesting that variability within the diseased state may be related to disease progression due to the effects of the treatment.

This study also analyzed the presence of differentially abundant proteins (DAPs) in OA across various treatment groups, including YG versus Normal Control (YvN), diclofenac versus Normal Control (SvN), Disease versus Normal Control (DvN) and Vehicle versus Normal Control (VvN) (Table S6, Data S31 to S35).

To further investigate the effect of these DAPs, we conducted canonical Pathway analysis using IPA. The heat map (Figure 5, C) depicts pathways with an absolute z-score difference greater than two when comparing any conditions between YvN, DvN, and VvN. Each cell represents the canonical pathway z-score returned by IPA. Pathways with z scores greater than two are significantly activated. And pathways with z**-**scores less than negative two indicate inhibition. The heat map also shows that DvN forms a distinct cluster, characterized by strong activation of inflammatory and wound-response pathways and concurrent suppression of collagen/GP6 signaling. Both VvN, and more prominently, YvN converge toward the normal profile, dampening these OA-related inflammatory cascades while moderately restoring protein-handling and metabolic pathways. This aligns with findings that mitochondrial metabolism navigates inflammasome activation and bioenergetic control (Martínez-Reyes *et al*, 2020; Billingham *et al*, 2022; Mauro *et al*, 2011; Yoon *et al*, 2021).

### Synergistic Action of the Metabolites of YG

The synergistic action of YG’s bioavailable metabolites was studied by identifying overlapping drug targets with the TRGs of the OA DEGs undirected network. The interconnectedness of the PPI networks indicated that these metabolites have a synergistic action, which we further analyzed using the guilt-by-association approach. The basic principle in determining a synergistic action or partner is that two nodes of each type (metabolite/protein/pathway) should be connected through either a common or neighboring set of nodes of another kind (Figure 6, A) (Rai *et al*, 2021; Casas *et al*, 2019; Wildenhain *et al*, 2015). Therefore, to determine the synergy among YG metabolites, their target proteins, and associated pathways, we constructed a multi-component network comprising YG bioavailable metabolites, their potential protein targets (TRG of OA DEGs only undirected network), and associated pathways, or more precisely, the metabolite-target-pathway (MTP) network (Figure 6, B, Data S36). Path length threshold of ≤3 between pairs of bioavailable metabolites to identify interactions depicted in Figure 6, A, we created a metabolite–metabolite network, wherein nodes are synergistic couples if they share an edge (Figure 6, C).

**Figure 6:**
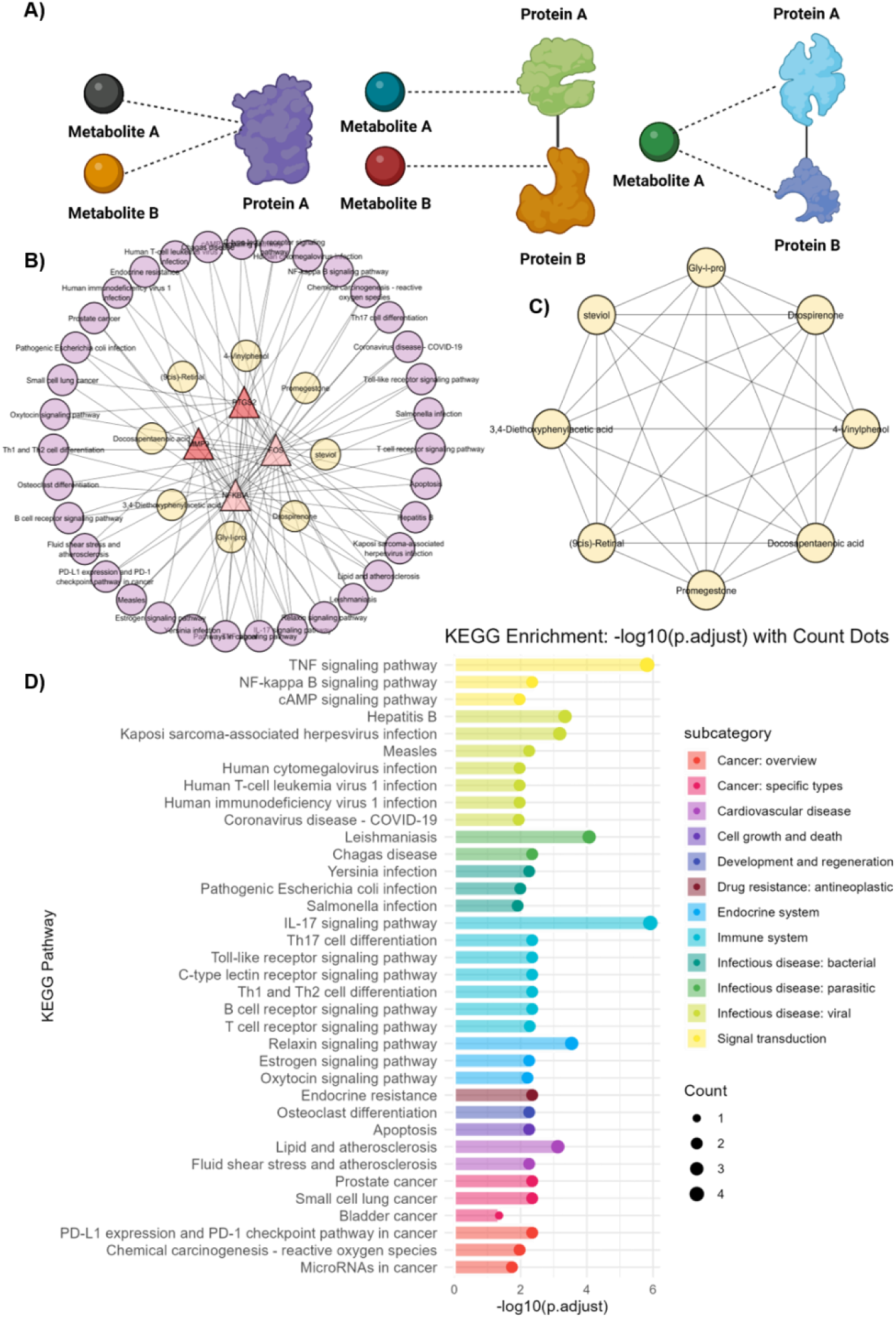
Synergistic interaction and network-based analysis of key therapeutic targets. (A) Representative protein–metabolite interaction models showing types of synergistic interaction between metabolites and proteins. Created in BioRender. (B) The molecular interaction network highlighting the OA DEG targets (triangles) connected to associated KEGG pathways (pink circles) and the instigating YG bioavailable metabolites (yellow circles). (C) The associated Metabolite synergistic network of YG bioavailable metabolites that target Key OA DEGs. (D) Comparative pathway enrichment analysis of the targets of bioavailable components of YG treatment conditions represented in the OA DEG network.

### Contextualizing the targets of YG in OA

Important determinants of the pathogenic pathways in OA and the cause for structural alteration of joints are NFKBIA, matrix metalloproteinase - 9 (MMP-9), prostaglandin-endoperoxide synthase 2 gene (PTGS2), and protein C-Fos. Notably, these four proteins also emerge as TRGs in our undirected, DEG-based human OA network. They are likewise targets of the exogenous DAMs found in rat plasma after YG administration. We focus on this high-confidence set to keep a balance between mechanistic insight and analytical clarity. The strict filter does leave out other possible candidates. However, this focus is necessary to unravel YG’s complex, multi-component mode of action without burdening the narrative with excessive constituent-associated details.

NFKBIA encodes IκBα, a pivotal inhibitor of NF-κB that sequesters NF-κB in the cytoplasm, thereby curtailing its ability to activate transcription of pro-inflammatory and catabolic genes. By directly inhibiting the dimeric NF-κB/REL complex, NFKBIA attenuates inflammatory responses (Hoffmann *et al*, 2002). While NF-κB normally helps to regulate cartilage growth and development, its abnormal, sustained activation in OA shifts its role from supportive to destructive, thereby fueling inflammation, promoting enzyme-mediated ECM degradation, and ultimately leading to joint degeneration (Tian *et al*, 2024; Mendez *et al*, 2020). Notably, aged articular cartilage exhibits a significant elevation in NF-κB activated chondrocytes, and enhanced IKK–NF-κB signaling accelerates age-related joint tissue degeneration (Catheline *et al*, 2021).

Furthermore, overexpression of NFKBIA in OA synovial fibroblasts effectively downregulates the expression of destructive enzymes, such as MMP-1, MMP-3, MMP-13, and ADAMTS4. This reduces inflammation and cartilage degradation. NF-κB’s role has been established in cellular energy metabolism and metabolic adaptation(Bondeson *et al*, 2007). These findings underscore the therapeutic potential of targeting NF-κB signaling in OA management.

MMP-9, known as gelatinase B, accentuates cartilage degradation by cleaving denatured collagens and aggrecans. Collagens and aggrecans are essential components that confer tensile and compressive strength to the articular cartilage(Slovacek *et al*, 2021). In the osteoarthritic milieu, increased MMP-9 expression facilitates the breakdown of extracellular matrix components and persists through its association with neutrophil gelatinase-associated lipocalin (NGAL), which safeguards its activity from auto-degradation. This sustained enzymatic function of MMP-9 amplifies cartilage destruction and joint deterioration(Meszaros & Malemud, 2012). Collectively, the orchestrated actions of collagenases and gelatinases, with MMP-1, exerting a predominant destructive influence and MMP-9 reinforcing matrix degradation, highlight the potential for targeted inhibition strategies in mitigating osteoarthritic progression(Mixon *et al*, 2021).

PTGS2 produces cyclooxygenase-2 (COX-2) enzyme, which converts arachidonic acid into prostaglandins to produce inflammation, pain, and cartilage destruction in OA. Normally unexpressed in most cells, COX-2 is robustly induced during inflammation, and its elevated expression in OA is associated with damaged articular cartilage and aberrant subchondral bone formation. Studies have shown that OA patients, along with rheumatoid arthritis patients, have elevated COX-2 levels, which lead to increased PGE2 production through osteocyte overexpression, which worsens joint degeneration(Chen *et al*, 2019; Fan *et al*, 2015). Preclinical investigations further reveal that genetic knockout of COX-2 in osteocytes or treatment with selective COX-2 inhibitors can rescue subchondral bone structure and attenuate cartilage destruction (Tu *et al*, 2019). These findings underscore the therapeutic potential of targeting PTGS2 through selective inhibitors, natural anti-inflammatory compounds, and gene regulatory therapies to modulate inflammatory cascades and mitigate joint degradation in OA.

Protein c-Fos, encoded by the FOS gene, is a proto-oncogene that forms an essential component of the AP-1 transcription factor complex through heterodimerization with Jun family members(Wagner & Eferl, 2005). AP-1, activated by inflammatory cytokines, growth factors, and mechanical or oxidative stress, plays a critical role in joint and cartilage physiology. In OA, c-Fos expression is notably elevated in cartilage, which modulates chondrocyte responses to microenvironmental stress. Genetic inactivation studies in experimental OA models reveal that c-Fos not only influences the expression of classical targets such as matrix-degrading enzymes but also orchestrates a metabolic shift in chondrocytes from aerobic glycolysis towards enhanced pyruvate oxidation and TCA cycle activity, mediated by key enzymes like pyruvate and lactate dehydrogenases. These findings underscore the pivotal role of c-Fos in regulating cartilage integrity and energy metabolism, highlighting its potential as a therapeutic target in OA and possibly in other conditions marked by aberrant chondrocyte proliferation and metabolism.

### Proposed Mechanism of Action

Through multiple mechanisms, YG exerts significant therapeutic effects against OA. A reduction of inflammation occurred in the knee joint (according to H&E staining). The cartilage matrix and chondrocytes of the knee joint were preserved (according to Fast-Green Safranin staining). YG effectively stabilizes subchondral bone architecture of the knee joint (according to micro-CT imaging). Also, it alleviated pain symptoms and improved mobility (according to behavioral studies). The multi-component nature of YG allows simultaneous targeting of inflammation (NF-κB, TNF signaling, cAMP signaling, and other pathways associated typically with immune system activation due to infection), cartilage degradation (Collagen modulation, Extracellular matrix organization, relaxin signaling), and bone remodeling (Osteoclast differentiation) to manage OA (Figure 6, D, Data S37). We acknowledge that more targets are possible, which we have marked in the extended networks (Figure S7). However, increasing targets also increases complexity and the possibility of false attribution. These findings collectively suggest that YG targets proteins pivotal to these pathways, orchestrating a multi-pronged regulatory effect to counteract the pathological hallmarks of OA.

### Proteomic Evaluation of YG’s Proposed Mechanistic Effects against OA

The restorative effect of YG on MIA-induced OA in rats was assessed by comparing the DAPs of proteins in the disease versus normal (DvN) and YG-treated versus normal (YvN) groups. Proteins differentially abundant due to OA induction were identified in the DvN comparison, while changes in DAPs between the DvN and YvN groups were attributed to treatment with YG. The primary distinction between these groups was the treatment intervention, and therefore any observed shifts in protein abundance were considered treatment-related effects.

Proteins were categorized based on their differential abundance in disease and YG treatment compared to normal animals (Table S7). The following categorization arose.

- **No effect of Yograj Guggulu DAPs.** Proteins identified as DAPs under DvN and YvN conditions indicate that the treatment did not significantly modulate them. A total of 430 DAPs were categorized as not effected by YG treatment.
- **Normalized Proteins: Disease-associated DAPs.** Proteins identified as DAPs in DvN but not in YvN suggest restoration of normal abundance due to the treatment with YG. The 364 DAPs can be categorized as normalized proteins (Data S38).
- **Treatment-specific DAPs.** Proteins identified as DAPs exclusively in YvN suggest treatment-related modulation independent of the disease state. A total of 71 DAPs can be categorized as treatment-specific DAPs.

This categorization provides a systematic approach to evaluate proteomic changes associated with OA induction and the potential therapeutic effects of YG. The subset of proteins uniquely dysregulated in untreated MIA-induced OA (DvN) but not in YG-treated animals (YvN) mapped overwhelmingly to innate-immune and antigen-presentation pathways. They also featured prominently in central carbon and nucleotide metabolism as well as the biosynthesis of essential cellular building blocks. Beyond metabolism, these proteins drove protein processing in the endoplasmic reticulum, mediated nucleocytoplasmic transport, organized the cytoskeleton and focal adhesions, and contributed to broader systemic regulatory networks. Their return to non-differential abundance in YvN demonstrates that YG specifically reverses these OA-related molecular perturbations, such as dampening synovial inflammation, restoring chondrocyte bioenergetics and biosynthesis, alleviating ER stress, preserving matrix integrity and cell-matrix adhesion, and normalizing local RAS/calcium signaling (Figure 7, Data S39). This pathway-level normalization mirrors the histological preservation of cartilage, micro-CT stabilization of subchondral bone, and behavioral pain relief observed after treatment.

**Figure 7:**
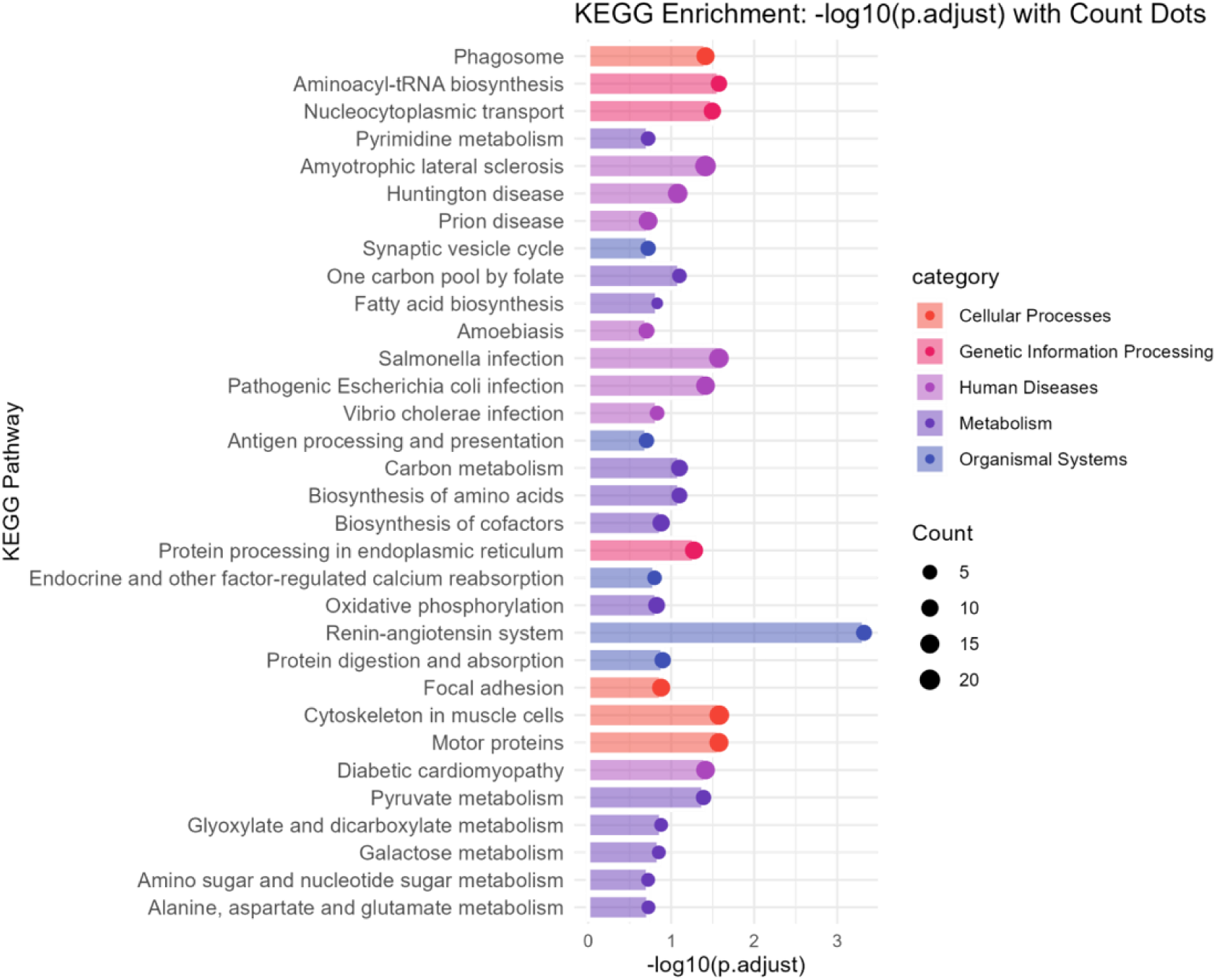
YG Normalises the MIA-induced OA DAPs. Bar charts depicting top functional pathways and biological processes enriched in each treatment DAPs that normalized, highlighting changes in cartilage matrix organization, inflammatory response, and other key processes underlying joint pathology. Table S7 summarizes the DAPs that remain unaffected or normalized due to the treatment when compared to disease DAPs with normal control group expression as baseline.

### Shortest Path Analysis of YG

Following treatment with YG, a subset of these DAPs returned to near-baseline expression levels (in YvN comparisons), suggesting a restorative effect of the treatments. To explore the potential regulatory relationship between the inferred drug targets of YG and these normalized proteins (Figure S8, A), we first performed pathway enrichment analyses on drug targets identified as TRGs within an undirected network constructed solely from DEGs. Although shared pathway involvement suggests possible functional relationships, pathway enrichment alone does not establish directional or mechanistic causality (Yuan *et al*, 2021; Gerassy-Vainberg *et al*, 2024; Castel *et al*, 2016). Therefore, we conducted shortest path analysis on a directed extended network, setting YG drug targets as source nodes and the normalized proteins as target nodes. Shortest path analysis allowed us to trace potential regulatory cascades, providing mechanistic evidence by identifying the most direct and biologically plausible signaling routes connecting the drug targets to the normalized proteins. Short, directed paths supported the hypothesis that the therapeutic effects of YG are mediated through the modulation of specific molecular intermediates, thus reinforcing a causal rather than merely correlative interpretation of our findings. We identified the shortest path to 45 of the 70 normalized proteins, within the KEGG pathways associated with YG’s targeting (Figure S 8, B, Table S8, Data S40).

## Discussion

Osteoarthritis (OA) is a chronic and widespread disease with societal impact. Conventional treatment of OA is inadequate, and currently, there are no approved disease-modifying drugs for osteoarthritis (DMOADs). Recent serum molecular biomarkers demonstrate that knee OA can be predicted up to eight years before x-ray abnormalities appear (Kraus *et al*, 2024). This broadens the temporal window for intervention and highlights the systemic nature of OA. We explored the effects and mechanisms of Yograj Guggulu (YG), a traditional Ayurvedic formulation, in OA. Using an integrated approach combining transcriptomics, metabolomics, proteomics, and advanced network analysis, we gained significant insights into the broad disease-modifying effects and the biological pathways modulated by YG. Ayurveda is an ancient system of medicine that originated more than 3000 years ago. It is known for its holistic approach to overall health and well-being. The term ‘Ayurveda’ combines two words ‘Ayuh’ which means ‘life’ and ‘veda’ means ‘knowledge’. Ayurveda treats the system as a whole, and do not specifically address isolated symptoms, which makes it a classic case for a network medicine study (Valiathan, 2009; Gan *et al*, 2023).

Distance-based network analysis of differentially expressed genes enabled metformin repurposing for atrial fibrillation(Lal *et al*, 2022). Similarly, utilizing the knowledge on differentially expressed genes in OA from the integrated transcriptome of human synovium, we constructed a network and paired it with bioavailable metabolites of YG (in rats) to find their core targets in OA. Using pathway enrichment analysis, we identified significant overlaps between the biological processes enriched pathways, potentially targeted by YG, and those disrupted in OA. The key pathways, such as the NF-κB signaling, IL-17 signaling, collagen degradation, and osteoclast differentiation with important roles in inflammatory regulation and cartilage homeostasis, were identified. We also predicted that the bioavailable metabolites worked in synergy, based on the closeness centrality of the metabolite–metabolite network. This synergy within the metabolites implied that the compounds act in concert rather than targeting individual pathogenic nodes. In essence, our network model predicts that YG broadly counteract the molecular dysfunction of OA by modulating multiple disease-related pathways.

Rats treated with YG exhibited a significant reduction in the joint inflammation as was evident from micro-CT imaging and histology. Interestingly, preservation of the subchondral bone architecture, cartilage matrix, and chondrocytes were also observed. Comprehensive behavioral tests revealed that YG-treated rats experienced reduced pain and improved locomotor function. The protective effect of YG was comparable to that of diclofenac sodium. Notably, YG-treated rats had no significant damage to the stomach mucosa, which presents a distinct advantage compared to diclofenac sodium treatment.

We observed that the expression of several proteins, which were altered in the joint tissues on OA induction, reverted to near-normal levels upon YG administration. These proteins included MMPs, NF-κB pathway components, and inflammatory mediators, all known to be involved in the pathogenesis of OA. Restoration of pathways related to immune response, metabolism of biomolecules, energetics, and matrix homeostasis links proteomic normalization, with the observed improvement in the structure and function of OA-affected joint.

Our shortest path analysis on the directed extended network of the disease provided additional context to the causal relationships between drug targets and normalized proteins. This analysis enhanced the interpretative power of our results beyond mere correlation by tracing regulatory cascades from drug targets to the proteins modulated by the treatment. Specifically, we could determine the chain of protein cascades from the targets of YG to the 45 normalized proteins that were also part of the pathways enriched within the targets. This enabled us to determine the modulation of NF-κB signaling, IL-17 signaling, collagen degradation, and osteoclast differentiation pathways with important roles in inflammatory regulation and cartilage homeostasis. By finding this causal interaction, we could validate the multitarget multi-component interaction of YG to ameliorate OA systemically. This analysis further revealed specific molecular interactions, identifying regulatory targets that could be explored for future mechanistic studies or targeted interventions.

A limitation of our study is that our findings require confirmation of the predicted interactions through targeted assays or orthogonal methodologies. Our metabolomic analyses revealed many bioactive metabolites. However, batch-to-batch variability within a threshold cannot be completely ruled out in the case of herbal formulations, and hence, independent verification is desirable. For clinical use, combinations of bioavailable chemical constituents are to be considered. The similar framework has shown translational success around single pathways (Gerassy-Vainberg *et al*, 2024).

In conclusion, our integrated omics approach and advanced network analysis demonstrate significant potential for YG in the treatment of OA. Our study also reinforces the value of an approach of employing modern scientific validation and robust analytical framework for identifying and validating therapeutic targets of traditional medicines for the management of chronic complex diseases. Collectively, our findings also exemplify how Ayurvedic polyherbal therapy can be rationalised within a contemporary network-medicine framework. Thereby offering a data-driven blueprint for holistic yet mechanism-rich drug discovery.

## Methods

### Large-scale expression analysis

The large-scale expression analysis was performed using MIAME-compliant microarray datasets obtained from the NCBI GEO database. Three Affymetrix platforms were used, namely, the Human Genome U133A (GPL96), Human Genome U133B 2.0 (GPL97), and Human Genome U133 Plus 2.0 (GPL570) arrays. Only studies utilizing fresh or flash-frozen human tissue samples with at least four samples per study were selected. Raw data were downloaded using the getGEOSuppFiles function from the GEOquery R package. For each platform, tar files were extracted and .CEL files were read using the ReadAffy function from the affy package. Data normalization was carried out using fRMA (with hgu133afrmavecs or hgu133plusfrmavecs, depending on the microarray platform) or the Robust Multiarray Average (RMA) method, which included background correction and log2 transformation.

Following normalization, datasets were integrated by merging on common probes. Dimensionality reduction using Uniform Manifold Approximation and Projection (UMAP) from the umap package (McInnes *et al*, 2018) enabled tracking of batch effects, which were subsequently corrected with ComBat from the sva package (Leek & Storey, 2007; Leek *et al*, 2019). The limma package was used for a differential expression analysis comparing OA and normal groups. Genes were considered differentially expressed if they exhibited an absolute log fold change of ≥1.0 and a False Discovery Rate (FDR) below 0.01 (or below 0.05 for the GPL97 dataset). Differentially expressed genes across the datasets were then used as seeds for subsequent disease network construction.

### Network construction

Protein-protein interactions (PPI) were retrieved from the STRING database (Szklarczyk *et al*, 2023) to construct the undirected interaction network using high-confidence interactions with a score of 400. The statistical significance of the interactions was assessed using p-values provided by STRING, which are calculated using a hypergeometric test. The interactions were then reconstituted into a binary adjacency matrix (A), where an interaction between two proteins was denoted as 1, and no interaction was denoted as 0. For the construction of the directed network, we employed NetControl4BioMed(Popescu *et al*, 2021). This tool was used to establish the protein-protein interaction (PPI) using data from Omnipath(Türei *et al*, 2016), InnateDB(Breuer *et al*, 2013), SIGNOR(Licata *et al*, 2020), and Human Interactome databases, and directionality was assigned to the interactions based on the regulatory relationships between the nodes. Using the input data, a directed adjacency matrix was generated, where a directed interaction between two proteins was denoted as 1 and 0 if no interaction was present. Subsequently, key topological features of the network, including structural properties such as degree, clustering coefficient, betweenness centrality, and network diameter, were computed to analyze the overall network structure.

In the extended network, the neighbors (interaction partners) of the mapped proteins were also retrieved, expanding the network to include proteins not directly found in the initial DEGs list but related through their interactions using the “neighbors” function from the STRING R package for the undirected network and NetControl4BioMed web tool for the directed network.

### Topologically relevant gene identification

The largest connected component was isolated for each network, and key topological parameters, degree and betweenness centrality, were computed using Cytoscape’s (Shannon *et al*, 2003) network analysis suite. Nodes with betweenness centrality values in the top 80th percentile were classified as bottlenecks, signifying their critical role in mediating communication across the network. Independently, nodes with degree values in the top 80th percentile were identified as hubs, regardless of their betweenness centrality, reflecting their extensive connectivity within the network. Nodes exhibiting high betweenness centrality but lower degrees were categorized as non-hub bottlenecks, indicating their potential role in controlling network flow despite limited direct interactions.

### Experimental animal condition and ethical approval

The experimental protocols were approved by the Institutional Animal Ethics Committee (NIPER/PC/2022/46), and a study was conducted in accordance with the guidelines by CCSEA (Regd.No.: 2069/GO/Re/S/19/CPCSEA), India. Eighty Sprague Dawley rats (males only) at 8-9 weeks old were used for the experimental work. Up to 4 rats were housed per cage in a sanitary room with controlled temperature, humidity, and a 12-hour/12-hour light-dark cycle. Food and water were provided ad libitum.

OA was induced in the anesthetized rats by adopting the standard procedure with slight modifications. The rats received a single intra-articular injection of MIA. MIA was injected at 5mg/50µl, where 5mg MIA was dissolved in 50µl sterile saline. Under anesthesia, the left knee was shaved and disinfected, and then an incision was made at the center of the knee to denote the infrapatellar ligament. Each rat was positioned on its back, and the left leg was flexed 90° at the knee joint. MIA was injected into the medial side of the ligament of the left knee using a 29-gauge, 0.5-inch needle. Care was taken to ensure the needle did not reach too far into the cruciate ligaments.

### YG Dosing

Three YG dose groups were considered for the investigation. 135 mg/kg as low, 270 mg/kg as medium, and 540 mg/kg as high doses. The medium dose corresponds to the human equivalent dose of YG, approximately the therapeutic daily dose in a 70 kg adult. The low and high doses represented approximately half and twice this equivalent dose, respectively. All doses were administered once daily for 21 consecutive days by oral gavage. YG was prepared fresh each day by suspending the calculated dose in distilled water, and the suspension was delivered using an oral gavage tube. The control group received the vehicle (distilled water) alone under the same schedule.

### Behavioral Efficacy Assessments

To assess the phenotypic differences at the behavioral level, we conducted a series of behavioral tests at various time points (days 7, 21, and 28) during the treatment period. These tests included the Von Frey Hair Test, Open Field Test, Rotarod Test, Hot Plate Test, and Walking Track Analysis. The results provided insights into the analgesic and anti-inflammatory effects of the treatments, as well as their impact on locomotion and pain sensitivity. The body weight of the rats was also recorded every week (days 7, 14, 21, and 28).

### Von Frey Hair Test

For assessment of tactile allodynia, the hindlimb withdrawal threshold evoked by stimulation of the hind paw by von Frey filaments was determined while the rat was placed on a metal mesh floor with 96mm × 196mm x 140mm cells. After a 10-minute acclimation period, mechanical stimuli were applied to the left foot pad with 6 calibrated filaments (Ugo basile, Italy) ranging from 0.5 to 100.0 g until the filament bent. A single trial of stimuli consisted of 6 applications of filaments at a frequency of 1/second. Five trials were performed at 3-minute intervals.

### Open Field Test

The OFT apparatus (50×50×38 cm^3^) was divided into one center rectangular area and four corner squares. Initial training, which was 10 minutes, was conducted, followed by a five-minute test. Each rat was kept in a wooden box painted black from the inside. The open-field box was sanitized with 70% ethanol before spotting another animal. The OFT was performed on the 7^th^ day, 14^th^ Day and 28^th^ Day. Further, the distance traveled (m), immobility time (s), mobility time (s), and average speed (m/s) were recorded through a video camera, and tracking was plotted using Any-maze version 7.0 software.

### Rotarod Test

A rotarod test was done to check the performance of the hind limb, a motor coordination parameter. After initial training, rats were kept on a rotarod in an accelerating mode from 4 to 45 rpm with a cutoff time of 300 s. Falling time in seconds was recorded automatically by sensors attached to the platform.

### Hot Plate Test

Response latency was used to gauge pain sensitivity or analgesic effects in rats using the Hot Plate Test. This measure represents the time it takes for the subject to exhibit a response (like paw licking or jumping) upon being placed on a heated surface. The heated plate induces a thermal stimulus, and the latency to the first observable response provides valuable information about the subject’s pain threshold and nociceptive behavior. A shorter latency suggests heightened pain sensitivity, while an increased latency may indicate reduced nociception or the analgesic effects of a substance under study.

### Walking Track Analysis

Rat paws were painted with non-toxic ink, and rats were allowed to walk on a white paper sheet. The distance between two successive paw prints was calculated in cm.

### Bioavailable components of treatment in Blood Plasma

At the end of the animal study, blood was collected via cardiac puncture and transferred into EDTA-coated tubes. The tubes were centrifuged at 14,000 × g for 5 minutes at 4 °C to separate the plasma. Plasma samples stored at –80 °C were subsequently thawed on ice, vortexed, and centrifuged at 1,620 × g for 20 minutes at 4 °C. For metabolite extraction, 100 µL of each plasma sample was mixed with 400 µL of methanol and incubated at –20 °C for 1 hour. The samples were then vortexed for 1 minute and centrifuged again at 14,000 × g for 20 minutes at 4 °C. The resulting supernatant containing the metabolites was carefully transferred to fresh MCT vials and dried using a speed vacuum concentrator (Savant SPD1010, Thermo Fisher Scientific). Finally, the dried extracts were resuspended in 100 µL of 50:50 methanol: water containing 0.1% formic acid for LC-MS analysis.

Untargeted metabolomics analysis was conducted using a Vanquish UHPLC system (Thermo Fisher Scientific) coupled to an Orbitrap Eclipse Tribrid mass spectrometer (Thermo Fisher Scientific) via a heated electrospray ionization (H-ESI) source. Chromatographic separation was performed using a Waters reversed-phase column (2.1 × 150 mm, 1.8 µm particle size; #186003540), maintained at 45 °C in still-air mode. Mobile phase A consisted of 0.1% formic acid in water, and mobile phase B consisted of 0.1% formic acid in methanol. The method employed a 15-minute run time at a constant flow rate of 0.350 mL/min, using the following gradient: 0 min, 0.5% B; 5 min, 50% B; 6 min, 98% B; held until 12 min; returned to 0.5% B at 13 min; equilibrated at 0.5% B until 13.1 min; and stopped at 15 min. A 5 µL injection volume was used, with the autosampler maintained at 5 °C. A diverter valve directed the flow to the mass spectrometer from 0.0 to 14.8 min and to waste thereafter.

Mass spectrometric analysis was conducted on the Orbitrap Eclipse Tribrid in Full MS/dd-MS² data-dependent acquisition (DDA) mode, operated in both positive and negative ion modes over the 15-minute run. The spray voltage was set to +3400 V (positive mode) and – 2800 V (negative mode), with a capillary temperature of 300 °C and vaporizer temperature of 400 °C. Sheath, auxiliary, and sweep gas flows were set to 40, 5, and 0 arbitrary units, respectively. The S-lens RF level was 55%. For MS¹ scans, data were acquired in the Orbitrap with a resolution of 120,000, an automatic gain control (AGC) target set to standard, and a mass range of m/z 100–1000. Data were recorded in profile mode with one microscan per scan. For MS² scans, data-dependent fragmentation was triggered on precursor ions exceeding an intensity threshold of 2 × 10⁴. Selected precursors were isolated using a 1.5 m/z window, fragmented by stepped higher-energy collisional dissociation (HCD) with collision energies of 20%, 35%, and 50%, and analyzed in the Orbitrap at a resolution of 15,000. MS² spectra were acquired in profile mode, with a maximum injection time of 22 ms, and a custom AGC target. Dynamic exclusion was enabled for 2.5 seconds, and isotope exclusion was applied to prevent repeated fragmentation of isotopic peaks.

20 mg of YG were weighed and dissolved in 1 ml of 80 % chilled methanol with 0.1 % formic acid. The mixture was vortexed for 10 minutes and incubated on ice for 10 minutes. This was followed by sonication for 10 minutes and incubation on ice for another 10 minutes. Subsequently, the mixture was centrifuged at 16,100 x g for 10 minutes at 4°C. An aliquot of 800 µl of the supernatant was drawn into a syringe and passed through a 0.22 µm filter to remove particulates.

### Feature annotation of medicinal extract molecular spectra by Compound Discoverer

Data were processed using Compound Discoverer software (version 3.3, Thermo Fisher Scientific). Data were processed using the built-in workflow template titled “Untargeted Metabolomics with Statistics: Detect Unknowns with ID using Online Databases and mzLogic.” The workflow comprised the following key steps: (1) retention time alignment using the ChromAlign algorithm, (2) compound filtering based on an Original Peak Rating threshold of ≥ 4, (3) compound grouping using a mass tolerance of 5 ppm and retention time (RT) tolerance of 0.2 minutes, (4) signal-to-noise filtering with a threshold of 1.5, (5) elemental composition prediction and gap filling, (6) compound identification using mzCloud and ChemSpider, (7) and similarity searching via mzLogic. Compound identification was further supported by cross-referencing multiple public databases, including BioCyc, KEGG, LIPID MAPS, NIST, NIST Chemistry WebBook, and NIST Spectra.

### Feature annotation of medicinal extract molecular spectra by Mzmine

To process the LC-MS data, the following methodology was applied using Mzmine(Schmid *et al*, 2023). First, the raw data files were imported, including Blank and YG samples in both positive and negative ionization modes. All MS levels were included, and no specific filtering was applied based on retention time, mobility, or scan number. The scan polarity was set to “any,” and the spectral library used for matching was loaded from a pre-defined location. Following data import, mass detection was performed across all files, including both Blank and Drug samples, using an automatic mass detector with a noise level threshold of 10,000 to eliminate low-intensity signals. Next, chromatogram building was conducted using the MS1 level. Chromatograms were constructed with a minimum of five consecutive scans, with an m/z tolerance of 0.002 m/z or 5.0 ppm. A minimum absolute peak height of 10,000 was set to filter out low-intensity peaks. Chromatograms were allowed to include single-scan features, and results were saved with a suffix indicating positive or negative mode chromatograms. The local minimum feature resolver was then applied to detect distinct features in the chromatograms. This step used a chromatographic threshold of 0.85 and a minimum search range of 0.05 for retention time. The MS1 to MS2 precursor matching was set with a tolerance of 0.01 m/z or 10 ppm, and the process was carried out for both positive and negative modes. Following this, the 13C isotope filter was used to group isotopic peaks, with a maximum charge state of 2. Features were deisotoped, ensuring that MS2-associated features were not removed. The final feature lists, both isotopic and deisotoped, were saved with a suffix to indicate the resolved and deisotoped nature of the chromatograms. Finally, spectral library matching was performed using a weighted cosine similarity approach. A precursor m/z tolerance of 0.001 m/z or 5.0 ppm and a spectral m/z tolerance of 0.0015 m/z or 10.0 ppm were employed. A minimum of four matched signals and a cosine similarity threshold of 0.7 were required for a successful match.

### Identification of compound targets

The protein targets associated with these drugs (similar to bioavailable YG chemical constituents) were considered the representative set of proteins with which the treatment’s chemical constituents can interact. This repositioning was based on the principle that structurally similar molecules often share targets. To identify potential drug targets for the treatment’s chemical constituents, we compared the structure of the treatment’s bioavailable chemical constituents with the structures of well-known modern drug molecules available in the DrugBank database(Knox *et al*, 2024). The similarity index, namely the Tc between drug molecules and metabolites, was generated using Open Babel(O’Boyle *et al*, 2011). Tc is defined as the ratio of the intersecting set of a molecule to the union set calculated as the similarity measure. Mathematically, Tc can be represented as Tc (a, b) = N_c_/ (N_a_ + N_b_ − N_c_), where N is the number of attributes in each molecule (a, b) and C is the intersection set.

### Determination of synergistic partners and their role

Using drug combinations against multiple linked targets in a disease-specific network is important for treating systemic diseases such as OA. Ayurveda is the science of combinatorial therapy, and its medicines often have multiple constituents, which are considered to operate through a cooperative mechanism termed synergy. Most of the research on combinatorial medicines has been based on the hypothesis that the synergistic action of drugs is due to their combinatorial effects on targets identified by their network topological features. Thus, we categorized the proteins in accordance with their corresponding metabolites and observed the presence of proteins and their corresponding metabolites and pathways in the maintenance of network integration. We use the guilt-by-association approach where the distance of ≤ three among the connected entities was considered for each dataset in undirected networks.

### Shortest-path analysis

The OA disease directed extended network was imported and annotated into R using the igraph package (Csardi & Nepusz, 2006; Csárdi *et al*, 2025). The shortest paths were determined from bioavailable YG TRG targets to the 70 normalized proteins.

These normalized proteins were those with abundance normalized after treatment and that share KEGG pathways with to YG TRG targets.

### Micro-CT

In vivo, micro-CT imaging was performed in Quantum GX2 Micro-CT (PerkinElmer). A quantitative analysis of the hind knee joints was performed using the micro-CT system. The specimens were scanned with an X-ray source of 90 kV, a field of view (FOV) of 36 mm, a pixel size of 88 µm, and a Cu 0.06 + Al 0.5 filter. The images were acquired in high-resolution mode; each scan took 14 minutes. The scan was acquired with a complete rotation of 360°. After scanning, we made both 3D reconstructions and calculations of osteogenesis properties. The volume of interest (VOI) was defined at the tibia metaphysis. The following parameters were calculated: bone volume (BV), cortex volume (CV), trabeculae volume (TV), trabecular tissue volume, whole volume, percent bone volume (BV/TV) and trabecular number (Tb. N). The reconstruction of the images was performed using Analyze 14.0 software, with the help of an add-on, bone microarchitecture analysis (BMD).

### Histopathology

After behavioral testing, the animals received ether anesthesia before their immediate decapitation, followed by fast removal of their left hind knee joint and stomach. Half of the rats’ left hind knee joints underwent storage in – 80° C using ice-cold saline solution while the other half were preserved in 10% formalin for histopathological examination.

The specimens containing knee joints from the tibia to the distal metatarsal with tarsal joint received 48 hours of formalin fixation at room temperature. The specimens underwent decalcification treatment for two weeks before they received paraffin embedding. The tissue specimens obtained from paraffin blocks underwent sectioning at 5 mm thickness before deparaffinization and rehydration through xylene, followed by absolute to 50% alcohol solutions. The articular tissue morphological changes became visible through H&E staining of the rehydrated sections.

At room temperature, the stomach received 48 hours of fixation in a 10% neutral-buffered formalin solution. The specimens underwent paraffin embedding after fixation. The paraffin blocks enabled the preparation of 5 µm thick sections, which underwent xylene deparaffinization followed by ethanol rehydration from 100% to 50%. The H&E staining procedure was applied to rehydrated sections to evaluate gastric tissue morphological changes.

### Tissue processing, protein extraction, and digestion of knee joint

Rat knee joints were dissected, lyophilized for 24 h, and pulverized in 2 mL tubes containing 1.4 mm acid-washed zirconia beads using a bead beater (1200 GenoLyte, SPEX SamplePrep). The powder was resuspended in 50 mM ammonium bicarbonate (ABC), vortexed briefly, pulverized again, and centrifuged at 13,000 g for 5 min at 4 °C to remove the beads. Guanidine extraction buffer (4 M guanidine-HCl, 50 mM sodium acetate, Roche Cocktail, and DTT 0.1 M) was made up in each sample vial such that the final volume remaining was 800 µL and incubated for 24 hours in a cold room on an orbital shaker.

The samples were centrifuged at 13,000 × g for 30 minutes at 4 °C, and the supernatant was collected. To the remaining pellet, 200 µL of 0.1% RapiGest extraction buffer (Waters) prepared in 50 mM ammonium bicarbonate (ABC) was added, followed by brief vortexing.

The mixture was centrifuged again at 13,000 × g for 30 minutes at 4 °C, and the resulting supernatant was collected. Protein-containing supernatants were desalted using 3 kDa molecular weight cut-off (MWCO) filters (Amicon® Ultra-0.5 Centrifugal Filter Units, Millipore), and protein concentrations were determined using the BCA assay. A total of 100 µg of protein from each sample was digested with trypsin (0.4 µg/µL), and the resulting peptides were purified using C18 columns (Pierce™ C18 Spin Tips, Thermo Fisher Scientific). After speed-vacuum drying, the peptides were reconstituted in 100 µL of 0.1% formic acid prior to LC-MS analysis.

### Liquid chromatography conditions for proteomics analysis

Proteomic analysis was performed using a Dionex UltiMate 3000 HPLC system (Thermo Fisher Scientific) coupled to an Orbitrap Eclipse Tribrid mass spectrometer (Thermo Fisher Scientific). Approximately 1 µg of peptides was injected and separated on an EASY-Spray PepMap RSLC C18 column (75 µm × 500 mm, 2 µm particle size, 100 Å pore size; Thermo Fisher Scientific) maintained at 40 °C. The mobile phases consisted of Solvent A (0.1% formic acid in ultrapure water) and Solvent B (0.1% formic acid in 80% acetonitrile). The injection volume was 2 µL, and the flow rate was set at 0.250 µL/min. The total run time was 145 minutes.

The system was equilibrated at 5% Solvent B for 10 minutes prior to sample injection. Following injection, the gradient was held at 5% B from 0 to 5 minutes. Solvent B was then gradually increased to 45% over the next 90 minutes (5 to 95 minutes), followed by an increase to 60% between 95 and 110 minutes. The percentage of Solvent B was further raised to 95% from 110 to 120 minutes and maintained at 95% for 10 minutes (120 to 130 minutes). The system was re-equilibrated by returning to 5% B over 1 minute (130 to 131 minutes) and held at this composition until the end of the run at 145 minutes. Gradient transitions were executed using a curve setting of 5.

### LC-MS/MS acquisition parameters for proteomics analysis

Peptide analysis was performed in positive ion mode using a nanospray ionization (NSI) source with a static spray voltage of 1400 V. The ion transfer tube was maintained at 275 °C. MS data were acquired in a data-dependent acquisition (DDA) mode with a cycle time of 3 seconds. Full MS scans were performed in the Orbitrap detector at a resolution of 120,000, over a mass range of m/z 350–1650, using quadrupole isolation. The AGC target was set to standard, with automatic injection time, and data were recorded in profile mode. Monoisotopic precursor selection was enabled for peptides, and only precursors with charge states from +2 to +7 and intensities above 1.0 × 10⁴ were selected for fragmentation. Dynamic exclusion was enabled to exclude a precursor after one selection for 6 seconds within a ±10 ppm window, with isotope exclusion turned on. For MS/MS acquisition, data-dependent higher-energy collisional dissociation (HCD) fragmentation was performed in the ion trap using a quadrupole isolation window of 1.6 m/z. The normalized HCD collision energy was set to 28%, and data were acquired in centroid mode at a rapid scan rate.

### Proteomics data processing parameters

Raw data files were analyzed using Proteome Discoverer version 2.5.0.400 (Thermo Fisher Scientific). Spectra were searched against the UniProt rat protein database retrieved on 24-01-05 using the Sequest HT search engine. The precursor mass tolerance was set to 20 ppm, and the fragment mass tolerance to 0.6 Da. Enzyme specificity was set to trypsin, allowing up to two missed cleavages, with a minimum peptide length of 6 and a maximum of 144 amino acids. Carbamidomethylation of cysteine (+57.021 Da) was specified as a static modification, while oxidation of methionine (+15.995 Da) and N-terminal acetylation (+42.011 Da) were set as dynamic modifications. Peptide identification was refined using the Percolator node, employing a target-decoy approach to control the false discovery rate (FDR). Strict and relaxed FDR thresholds were set at 1% and 5%, respectively. Peptides were further processed using the Minora Feature Detector node for label-free quantification, with a minimum trace length of 5, signal-to-noise threshold of 1, and a maximum isotope pattern retention time deviation of 0.2 minutes.

### Statistical analysis

All analyses were performed in R (v 4.4.2), GraphPad Prism 10 (GraphPad Software, San Diego, CA, USA) and Cytoscape (Shannon *et al*, 2003). Animals were randomly assigned to experimental groups. Each experimental group comprised eight male Sprague-Dawley rats. All eight animals per group underwent the battery of behavioural tests. After sacrifice, tissue allocation proceeded as follows: four randomly selected animals were processed for histopathological examination, and the remaining four were used for proteomic and plasma metabolomic profiling. From the histopathology set, three samples were randomly chosen for ex-vivo micro-CT imaging. Behavioural tests were analysed with two-way ANOVA (factors: treatment and disease followed by Tukey’s post hoc analysis. For omics data, biomolecules (genes, proteins, metabolites) were deemed differentially expressed when |log₂ fold-change| ≥ 1 with a Benjamini–Hochberg-adjusted p < 0.05. More stringent p-adjustment was adopted when the sample size was larger. Functional enrichment was evaluated using KEGG pathway analysis (threshold q < 0.05) and Ingenuity Pathway Analysis (IPA), considering pathways with |activation z| ≥ 2 as significantly perturbed.

## Data and code availability

The human osteoarthritic-synovial tissue transcriptome datasets analysed in this study are available from the NCBI Gene Expression Omnibus under accession numbers GSE12021, GSE55235, GSE55457, GSE55584, GSE12021, GSE206848, GSE236924, GSE43923, GSE82107, and GSE93698. The proteomics data have been deposited in the PRIDE repository (Perez-Riverol *et al*, 2016; Deutsch *et al*, 2023; Perez-Riverol *et al*, 2025), accession number PXD065118. Metabolomics data are accessible through MetaboLights (Yurekten *et al*, 2024), accession number REQ20250505210300. Micro-CT, histopathology datasets and the R code are available on Zenodo, 10.5281/zenodo.15728452. All other data supporting the findings of this work are contained within the Article and its Supplementary Information or can be obtained from the corresponding author upon reasonable request.

## Acknowledgement

The study was supported by IMR grant from the CCRAS, Ministry of AYUSH, Government of India, No. 3-27/2021-CCRAS. The study was supported by a PMRF research grant from the Ministry of Education, Government of India, to conduct this research. The proteomic/metabolic profiling was carried out utilizing liquid chromatography tandem mass spectrometry [LC–MS/MS] platform at the DBT-SAHAJ National Facility for Mass Spectrometry, BRIC-Rajiv Gandhi Centre for Biotechnology (RGCB), Thiruvananthapuram, Kerala. The authors thank RR histology services, Hyderabad, for assisting with the histopathology services of this study.

## Author information

## Contributions

VR and SP supervised the project and revised the manuscript. CCK and VR formulated the conceptual framework of this paper. AR performed most of the experiments, did network analysis and wrote the original draft. IB has done part of the network study. CCK, AJ and AS has revised the manuscript. BR, NP, and ANP conducted the behavioural experiments and collected the animal tissues. AR and DH assisted with the behavioural experiments. BR and AR performed the micro CT experiments. VGMN supervised the in vivo experiments. AJ, AR, NK, and AS conducted proteomics experiments. AJ, AR, and AS conducted metabolomics experiments. SP, NS, RA, GB, and SYR have contributed to designing the dosage of the medicine and made the ayurvedic medicine YG as per standard operating protocols.

## Declaration of interests

## Competing interests

This project was funded by CCRAS, New Delhi. The contents of the manuscript may have financial interests, especially pertaining to the therapeutic utility of YG constituents. IIT Guwahati has started the procedures for patenting the scientific content of this paper (Internal Reference Number: 2025063020000101).

